# Patterns of gene expression in pollen of cotton (*Gossypium hirsutum*) indicate down-regulation as a feature of thermotolerance

**DOI:** 10.1101/2021.06.06.447035

**Authors:** Farhad Masoomi-Aladizgeh, Matthew J. McKay, Yasmin Asar, Paul A. Haynes, Brian J. Atwell

## Abstract

Reproductive performance in plants is impaired as maximum temperatures consistently approach 40°C. However, the timing of heatwaves critically affects their impact. We studied the molecular responses of cotton male reproductive stages, to investigate the vulnerability of maturing pollen to high temperature. Tetrads, uninucleate and binucleate microspores, and mature pollen were subjected to SWATH-MS and RNA-seq analyses after exposure to 38/28°C (day/night) for 5 days. The results indicated that molecular signatures were down-regulated over developmental stages in response to heat. This was more evident in leaves where three-quarters of differentially changed proteins were decreased in abundance. Functional analysis showed that translation of genes increased in tetrads after exposure to heat; however, the reverse pattern was observed in mature pollen and leaves. Proteins involved in transport were highly abundant in tetrads, whereas in later stages of development and leaves, heat suppressed cell-to-cell communication. Moreover, a large number of heat shock proteins (HSPs) were identified in heat-affected tetrads, but these proteins were less abundant in mature pollen and leaves. We speculate that the sensitivity of tetrad cells to heat is related to increased activity of translation involved in non-essential pathways. Molecular signatures during pollen development after heatwaves provide markers for future genetic improvement.

## INTRODUCTION

An increase in global temperature is threatening food security by negatively affecting crop production across the world. The yields of crops are predicted to reduce substantially as global average temperature increases (Zhao et al., 2017); however, there is as yet no study showing the extent of yield reduction associated with the male gametophyte vulnerability to heat (Sita et al., 2017; Jiang et al., 2019; Djanaguiraman et al., 2018). Pollen development in flowering plants is composed of several stages that undergo meiotic and mitotic divisions to proceed to viable mature pollen for fertilisation (Borg et al., 2009), of which the early stage (post-meiotic tetrad cells) is extremely susceptible to high temperature (Begcy et al., 2019; Yu et al., 2019; Masoomi-Aladizgeh et al., 2020). Investigating molecular responses to extreme heat would help us understand the mechanisms of heat-sensitivity of the early stage of pollen development in plants.

Heat stress is initially sensed by the plasma membrane, which activates Ca^2+^ channels, causing an inward flux of Ca^2+^ into cells and activating the heat shock response (HSR) (Saidi et al., 2009). Mitogen activated protein kinases (MAPKs) and transcriptional regulators, such as heat shock transcription factors (HSFs) are activated, resulting in accumulation of stress-responsive proteins (Sita et al., 2017). Depending on the stage of pollen development at which plants experience heat, the thermotolerance is different, which can be due to distinct molecular responses (Masoomi-Aladizgeh et al., 2020). Previous studies have highlighted that the unfolded protein response (UPR) pathway in the endoplasmic reticulum (ER) and cytosol plays a crucial role in pollen response to heat stress. The ER and cytosolic UPR trigger bZIP transcription factors and HSFs, respectively, which are required to activate the HSR genes in the presence of unfolded proteins (Mittler et al., 2012; Raja et al., 2019). The function of the UPR is critical as the knocking out of *Inositol requiring enzyme 1 (IRE1)* in Arabidopsis, a component of the UPR pathway involved in the splicing of bZIP60, caused male sterility under heat stress (Deng et al., 2016).

Previous studies have highlighted that transcription and translation in plants undergo dramatic changes to mitigate the environmental stresses. It is known that alternative splicing events are induced when plants are subjected to environmental stresses. The post-transcriptional mechanism helps plant to adapt to harsh environments mainly by targeting the abscisic acid (ABA) pathway; however, defects in splicing machinery can lead to impairment in plant response to stress (Laloum et al., 2018). Moreover, Cui and Xiong, (2015) stated that alternative splicing (AS) of pre-mRNA occurs on a large number of genes when plant is exposed to abiotic stress. However, the increased m-RNAs under stress and limited splicing machinery, lead to splicing errors in heat-responsive genes. On the other hand, recent studies have demonstrated that down-regulation of non-essential pathways is undertaken by tolerant species to overcome stress conditions (Peredo and Cardon, 2020; Voesenek and Bailey-Serres, 2013). The new perspective on tolerance of plants proposes that up-regulation of protective genes must be warranted but the metabolism of cells should undergo an extensive down-regulation in response to abiotic stress, such as high temperature.

We aimed to study the transcriptome and proteome of four distinct stages of pollen development in cotton, in order to investigate molecular responses to 5 d of extreme heat (38°C). We hypothesised that the pollen developmental stage influences response to heat, and the early stage (tetrads) responds to heat differently. We predicted that tetrad cells would activate a large number of genes and proteins in response to extreme heat, compared with later developmental stages. We also hypothesized that down-regulation of functional pathways would be linked to thermotolerance of tissues and predicted that mature pollen and vegetative tissues would only up-regulate key heat-responsive functions.

## MATERIALS AND METHODS

### Plant growth and heat treatment

Sicot 71 seeds (*Gossypium hirsutum* L.) were germinated in plastic pots (21 × 26 cm; height × diameter) containing 9 kg soil mixed with roughly 18 g Nutricote controlled-release fertilizer (16 N:4.4 P:8.3 K). Plants were grown under glasshouse conditions at 30/22°C day/night temperature, and 12/12 h light/dark photoperiod under natural light. The light intensity during the day was maintained to a minimum of 400 µmol m^-2^ s^-1^ using supplemented light (Philips Contempa High Pressure Sodium lamps). Plants were fertilized weekly using Aquasol soluble fertiliser (Yates, Australia) at the rate of 4 g per 5 l and watered daily. Approximately 60 d after sowing, plants were acclimated to 34°C for 1 d, followed by exposure to 5 d extreme heat at 38/28°C. Vegetative and reproductive tissues including pollen, anther and leaf were all sampled. The glasshouse experiment was performed twice to collect adequate samples from different stages of pollen development for downstream analysis.

### Pollen collection and viability test

Squares corresponding to four distinct reproductive stages including tetrads (TE, 5.5-6 mm), uninucleate (UN, 6.5-10 mm) and binucleate (BN, > 11 mm) microspores and dehisced pollen (AN, anthesis) were collected according to the size of flower bud as described by Song et al., (2015). Sepals and petals were discarded using forceps, and the corresponding anthers were placed in 0.4 M mannitol solution. The mixture was vortexed gently to release pollen grains, followed by transferring the supernatant to a new tube. Pollen was centrifuged briefly, and the supernatant was discarded. The resultant samples were frozen in liquid nitrogen and stored at -80°C. To ensure that the collected pollen grains represented the specific stages of development, the nuclei were stained and visualised under an Olympus BX53 microscope as reported by (Masoomi-Aladizgeh et al., 2020).

To assess pollen viability, anthers corresponding to the four developmental stages were fixed in Carnoy’s solution (6:3:1 Ethanol: chloroform: acetic acid). The anthers were then placed on a microscopy slide and dissected to release pollen, followed by staining with 40 µL of Alexander’s solution (10 mL 95% EtOH, 1 mL malachite green (1% solution in 95% alcohol), 54.5 mL distilled water, 25 mL glycerol, 5 mL acid fuchsin (1% solution in water), 0.5 mL orange G (1% solution in water) and 4 mL acetic acid) according to Peterson et al. (2010). The slides were slightly heated and then observed under a Motic BA210 compound microscope.

### Thermal measurement and dehiscence

The surface temperature of reproductive and vegetative tissues was recorded with an infrared thermometer (Agri-Therm III, 6110L, USA). Under control and heat conditions, leaves and squares with a bloomed flower were selected, and their surface temperature was measured to compare the difference with the air temperature. For dehiscence, the heat-treated stages of pollen development (TE, UN, BN and AN) were allowed to recover under control conditions until they reached anthesis. Dehiscence corresponding to all developmental stages was then observed under a Motic SMZ-161 stereomicroscope.

### RNA isolation

Total RNA was isolated from approximately 15-50 mg of pollen developmental stages, including TE, UN, BN and AN, using a CTAB method optimized by Masoomi-Aladizgeh et al., (2016), followed by applying a RNeasy Plant Mini Kit (Qiagen). Briefly, a lysis buffer was prepared consisting of: CTAB (125 mg), EDTA (250 mg), Tris base (625 mg) and NaCl (1250 mg) in 25 ml nuclease-free water. The extraction buffer (500 μl) in addition to βME (25 μl) were added to the ground pollen, and the samples were incubated at 65 °C for 5 min. Chloroform (300 μl) was added to each tube, and the mixture was vortexed vigorously, followed by centrifugation at 14,000 rpm at 4 °C for 5 min. The supernatant (450 μl) was transferred into a QIAshredder Mini spin column to continue the isolation processes according to the manufacturer’s instructions. The concentration, purity and integrity of RNA was assessed with Epoch Take3 spectrophotometer (BioTek, Winooski, VT, USA) and TapeStation (Agilent Technologies, Santa Clara, CA, USA).

### Library preparation and Illumina sequencing

Library preparation and sequencing were conducted by the Ramaciotti Centre for Genomics (Sydney, NSW, Australia). QIAseq Stranded Total RNA library Kit (Qiagen, Germany) was used to convert RNA to Illumina sequencing libraries, following the manufacturer’s procedure. All 24 individual libraries were normalised and pooled together. The sequencing was then performed on a single flow cell lane of an Illumina NovaSeq 6000 system, generating 2 × 100 bp paired-end reads, with approximately 30 million reads per sample. A pass threshold of > 80% of bases higher than Q30 at 2 × 100 bp was used. FASTQ files were used for downstream analysis.

### Transcriptomics analysis

Paired-end FASTQ mRNA-seq reads were quality checked, trimmed, assembled and analysed using CLC Genomics Workbench ver 20.0.4.0 (Qiagen). Trimming was performed with the following parameters: Phred quality score >20 (limit=0.05); maximum number of two ambiguities; removing the terminal 15 nucleotides from 5’ end and 3 nucleotides from 3’ end; and removing truncated reads of <50 nucleotides length. Trimmed reads were then aligned to the cotton (*Gossypium hirsutum* L. acc. TM-1) reference genome (Zhang et al., 2015), with the following parameters: length fraction: 0.7; similarity fraction: 0.9; maximum number of hits for a read: 10. The transcript level for each gene was calculated by counting the number of reads mapped to each gene and normalised using the trimmed mean of M values (TMM) normalization method (Robinson and Oshlack, 2010). Differentially expressed genes were determined if the absolute fold change in expression was ≥3 with an FDR *p*-value of < 0.05 (Benjamini and Hochberg, 1995).

### Protein isolation and peptide preparation

Approximately 20 mg of pollen grains were ground into fine powder using a TissueLyser II (Qiagen), 5-6 Zirconox beads (2.8 to 3.3 mm) and liquid nitrogen. A phenol-based protocol was used for protein extraction. Briefly, 300 μL of protein extraction buffer (100 mM Tris-HCl, pH 8.0; 5% SDS; 10% glycerol; 10 mM DTT; 1% protease inhibitor) was mixed with the ground pollen grains, vortexed vigorously and incubated at room temperature for 30 min. The mixture was then incubated and shaken at 95°C for 3 min, followed by centrifugation at 20,000 × *g* for 10 min at 4°C. The supernatant (200 μL) was transferred into a new tube and mixed with an equal volume of 1.4 M sucrose, followed by extracting proteins twice with Tris-EDTA buffer-equilibrated phenol. Phenolic phase was carefully collected each time after centrifugation at 20,000 × *g* for 10 min at 4°C. The combined phases were again extracted using an equal volume of 0.7 M sucrose. Supernatants (300 μL) were transferred into a new tube, mixed with five volumes of cold 0.1 M ammonium-acetate in methanol and incubated at -20°C overnight. Samples were then centrifuged at 20,000 × *g* for 10 min at 4°C, followed by washing the pellets twice with cold 0.1 M ammonium-acetate in methanol and once with cold acetone. The pellets were finally air-dried and solubilized in 100 µL of 8 M urea in 100 mM Tris-HCl, pH 8.0. Protein concentration was measured by the Pierce BCA protein assay (Thermofisher Scientific, Waltham, USA).

Proteins from each replicate were digested in-solution with trypsin (Masoomi-Aladizgeh et al., 2020). Peptides were desalted using 47-mm SDB-RPS disks (3M, Saint Paul). The extraction disks were washed twice with 0.2% trifluoroacetic acid (TFA), followed by eluting the peptides in 200 μL of 80% acetonitrile (ACN) containing 5% ammonium hydroxide. Peptides were vacuum-dried and resuspended in 30 μL of 0.1% formic acid (FA). Micro BCA assay (Pierce, Thermo Fisher Scientific, San Jose) was used to quantify the peptides.

### High pH reversed-phase fractionation

A small fraction of all samples were pooled and fractionated using the Pierce High pH Reversed-Phase Peptide Fractionation Kit (Pierce, Thermo Fisher Scientific, San Jose). Briefly, 40 µg of the mixed peptides, from all samples analysed, was mixed and diluted with 300 μL of 0.1% TFA. The solution was loaded onto a high pH reversed-phase spin column, followed by centrifugation at 3000 × *g* for 2 min to retain the flow-through fraction. The same procedure was performed with Milli-Q water to retain an initial wash fraction. Eight gradient fractions (5%, 7.5%, 10%, 12.5%, 15%, 17.5%, 20% and 50%) were collected using the appropriate elution solutions. The samples (10 fractions) were vacuum-dried and reconstituted with 20 µL of 0.1% FA.

### Spectral library creation and SWATH-MS acquisition

Nanoflow LC-MS/MS was performed using a Triple TOF 6600 mass spectrometer (Sciex, USA) equipped with an Eksigent nanoLC 400 liquid chromatography system (Sciex, USA) and nanoflex cHiPLC module (Sciex, USA). For both data dependent and data independent acquisitions (DDA and DIA, respectively), the peptides were desalted with buffer A (2% acetonitrile with 0.1% formic acid) and loaded onto a C18 trap column (2 cm, 200 μm, 2.7 μm, Halo C18) at a flow rate of 5 µl.min^-1^ for 5 min. For DDA, the samples (10 µl, approximately 4 µg) were eluted from a cHiPLC C18 column (15 cm × 200 μm, 3 μm, ChromXP C18CL, 120 Å, 25°C, Sciex, USA) using a linear gradient of 2-35% of solvent B (90% acetonitrile with 0.1% formic acid) at a flow rate of 600 nL.min^-1^ in 120 min. For SWATH-MS (DIA) acquisition, the samples (10 µl, 2 µg) were separated in a linear gradient of 5-35% of solvent B at a flow rate of 600 nL.min^-1^ in 120 min. The peptides were then injected into a positive ion nanoflow electrospray MS analysis (spray voltage 2.5 kV, curtain gas 25, GS1 20) using a NanoSpray III source (Sciex, USA) and PicoTip Emitter (10 cm, 360 μm O.D., 20 μm I.D., New Objective, USA). To build a spectral library in DDA mode, high pH fractions were analysed using a TOF-MS survey scan over a mass range 350-1500 *m/z*, followed by 100-1800 *m/z* for MS/MS scans of the top 20 most intense precursor ions with an exclusion time of 30 s. To quantify the peptides in DIA mode, SWATH-MS was performed using a set of 100 variable *m/z* windows selected based on the intensity distribution of precursor *m/z* in DDA data. A TOF-MS survey scan was acquired (350-1500 *m/z*, 30 ms, high resolution setting of 30,000), followed by 100 SWATH-MS2 scans (350-1500 *m/z*, 30 ms, variable windows, high sensitivity setting of 15,000). SWATH-MS for each sample was acquired in a random order with one blank injection between samples.

### Proteomics analysis

Data generated in the DDA mode were processed using the Paragon algorithm (Sciex, USA) in ProteinPilot ver 5.0 (Sciex, USA) following modification of cysteine by carbamidomethylation, thorough ID mode, unused cut-off score of 1.3 (95% confidence) and false discovery rate (FDR) analysis. MS/MS spectra from the DDA run were searched against the *G. hirsutum* UniProtKB database released on September 2019 (78,943 sequences). ProteinPilot DDA search results were imported into PeakView (version 2.2, Sciex, USA) and used as a spectral library. To align retention times for all SWATH files, a linear regression was applied for each sample type by using at least five manually selected peptides across the elution profile. The following search parameters were used for quantitation: top 6 most intense fragment ions for each peptide, a maximum number of peptides of 100, 75 ppm mass tolerance, 99% peptide confidence threshold, 1% false discovery rate threshold and a 5 min retention time extraction window. The peak areas for peptides were extracted by summing the areas under curve (AUC) values of the corresponding fragment ions using PeakView. The summed peak areas of the peptides were log2 transformed and normalised, then used for protein quantification. To assess differentially expressed proteins, an analysis of variance (ANOVA) of the log-transformed normalised peak areas was carried out using two-sample student’s *t*-tests. Proteins with *p* < 0.05 and an expression fold-change greater than ± 1.5 with FDR < 0.05 by *t*-test were considered significantly changed between treatments (Wu et al., 2016).

### Gene ontology (GO) functional enrichment analysis

The resulting data set of differentially expressed genes (DEGs) and differentially expressed proteins (DEPs) were functionally annotated in OmicsBox (ver. 1.4.1). Fisher’s Exact Test was used to compare the lists of up- and down-regulated genes and proteins at each condition against the corresponding annotated database. Significantly enriched GOs were selected for further analysis (*p-value* < 0.05). The 20 most enriched GOs were retrieved for biological processes, and their semantic similarities were calculated using the SimRel approach implemented in REVIGO (Schlicker et al., 2006; Supek et al., 2011). The common enriched GOs between the tissues were selected, and their tendency to up- or down-regulation under heat stress was shown by polar plots in GOplot (Walter et al., 2015).

### Parallel Reaction Monitoring (PRM) Analysis

A total of 10 stress-responsive proteins identified and quantified by SWATH-MS were selected for targeted PRM quantification. Skyline (ver. 20.1.1.175) was used to create an inclusion list consisting of unique peptides with their mass to charge ratio (MacLean et al. 2010). Nanoflow LC-MS/MS using an Orbitrap mass spectrometer (Q Exactive, Thermo Fisher Scientific, Bremen, Germany) coupled to an Easy nLC 1000 nanoflow liquid chromatography system was operated in PRM mode (Hamzelou et al., 2020). Acquisition method included a single full MS scan combined with targeted MS/MS scans, with the following parameters: a resolution of 17,500, a target AGC value of 2 × 10^5^, a maximum injection time (IT) of 55 ms, and a normalized collision energy (NCE) of 30%). The PRM runs were analysed and quantified by Skyline using the sum of areas of all transition peaks for each peptide. The summed area was then log2 transformed, followed by *t*-test analysis.

## RESULTS

### Establishing a non-lethal elevated temperature

Our previous study showed that 40°C was lethal to the early stage of pollen development in cotton (Masoomi-Aladizgeh et al., 2020). Based on a pilot experiment, a daytime temperature regime was implied to induce a sub-lethal response to heat during microsporogenesis. Acclimating plants at 34°C for 1 d followed by exposure to 38°C for 5 d resulted in the development of mature pollen grains and therefore, samples that could be used for downstream molecular analysis. Figure 1A-B demonstrates the effect of 38°C on different stages of pollen development, resulting in damage specifically to the early stage of development (TE) compared with the later stages. The susceptibility of pollen to heat-induced damage was exacerbated by the consistently higher surface temperatures recorded at the surface of squares than on leaf surfaces when ambient temperatures were 38°C (Fig. 1C). For molecular analysis, square size and microscopy were used to identify each of the four pollen developmental stages (Fig. 2A). Chemical staining showed that pollen grains were viable in all stages of development after exposure to heat, enabling sampling and deeper molecular analysis. To analyse transcriptomes and proteomes at four distinct stages of pollen development under the elevated temperature, two independent glasshouse experiments, each with three replicates, were conducted to collect sufficient samples. RNA and protein isolation methods in this study were optimised for collection of very small amounts of pollen grains.

**Fig. 1.**
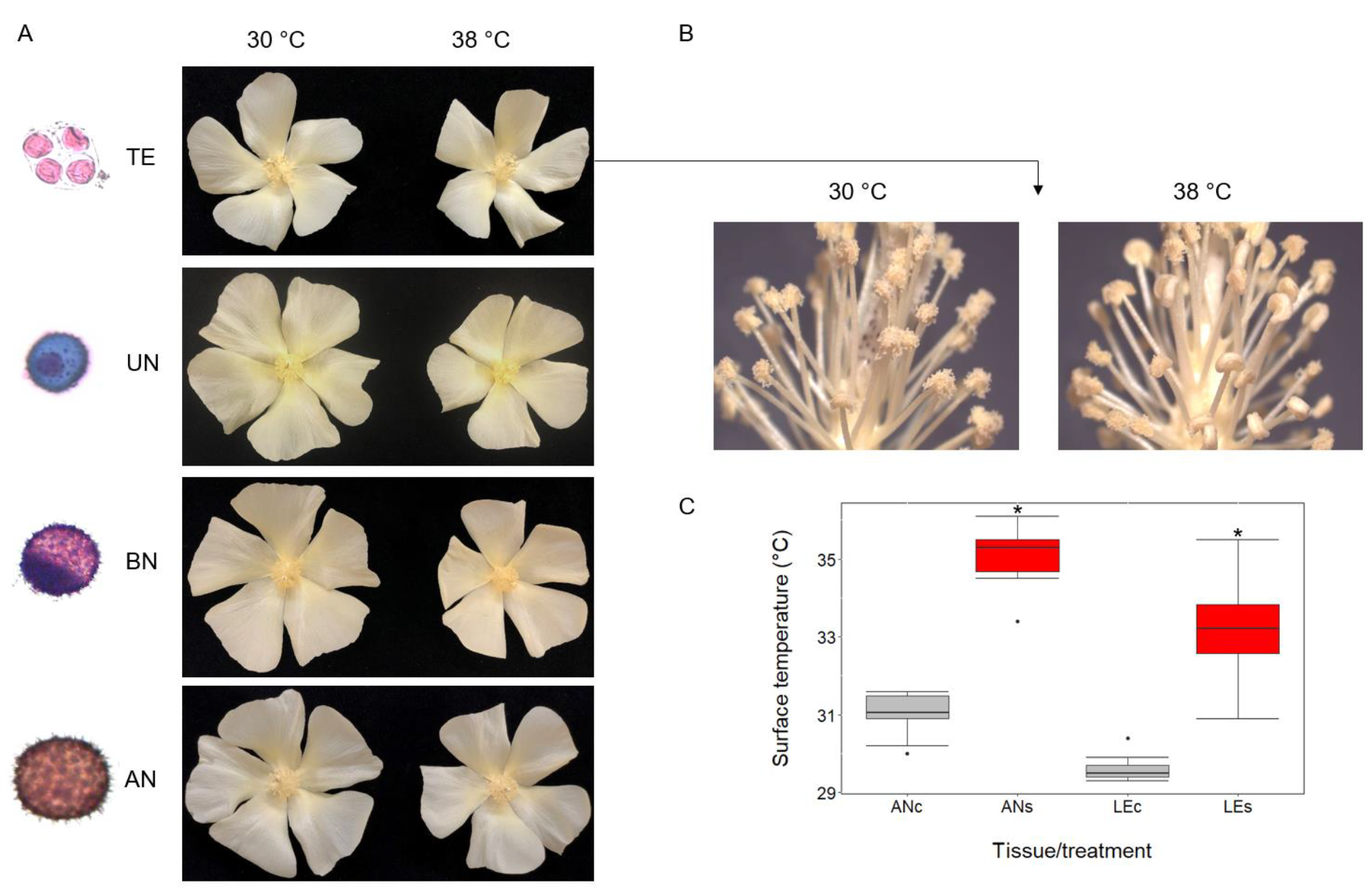
The effect of heat stress on pollen developmental stages in cotton. (A) the size of flowers at anthesis corresponding to TE, UN, BN and AN after exposure to 38/28°C for 5 d, compared with 30/22°C; (B) incomplete dehiscence of squares corresponding to the early stage (TE) exposed to 38/28°C for 5 d, compared with 30/22°C; (C) the real temperature of the squares at anthesis and leaves when exposed to extreme heat (ANc and ANs, squares at anthesis under control and stress conditions, respectively; LEc and LEs, leaves under control and stress conditions, respectively). The results are representative of at least 10 replicates. Asterisk (*) denotes a statistically significant difference between control and heat-stressed samples, as indicated by *t*-test analysis with *p*-value < 0.01.

**Fig. 2.**
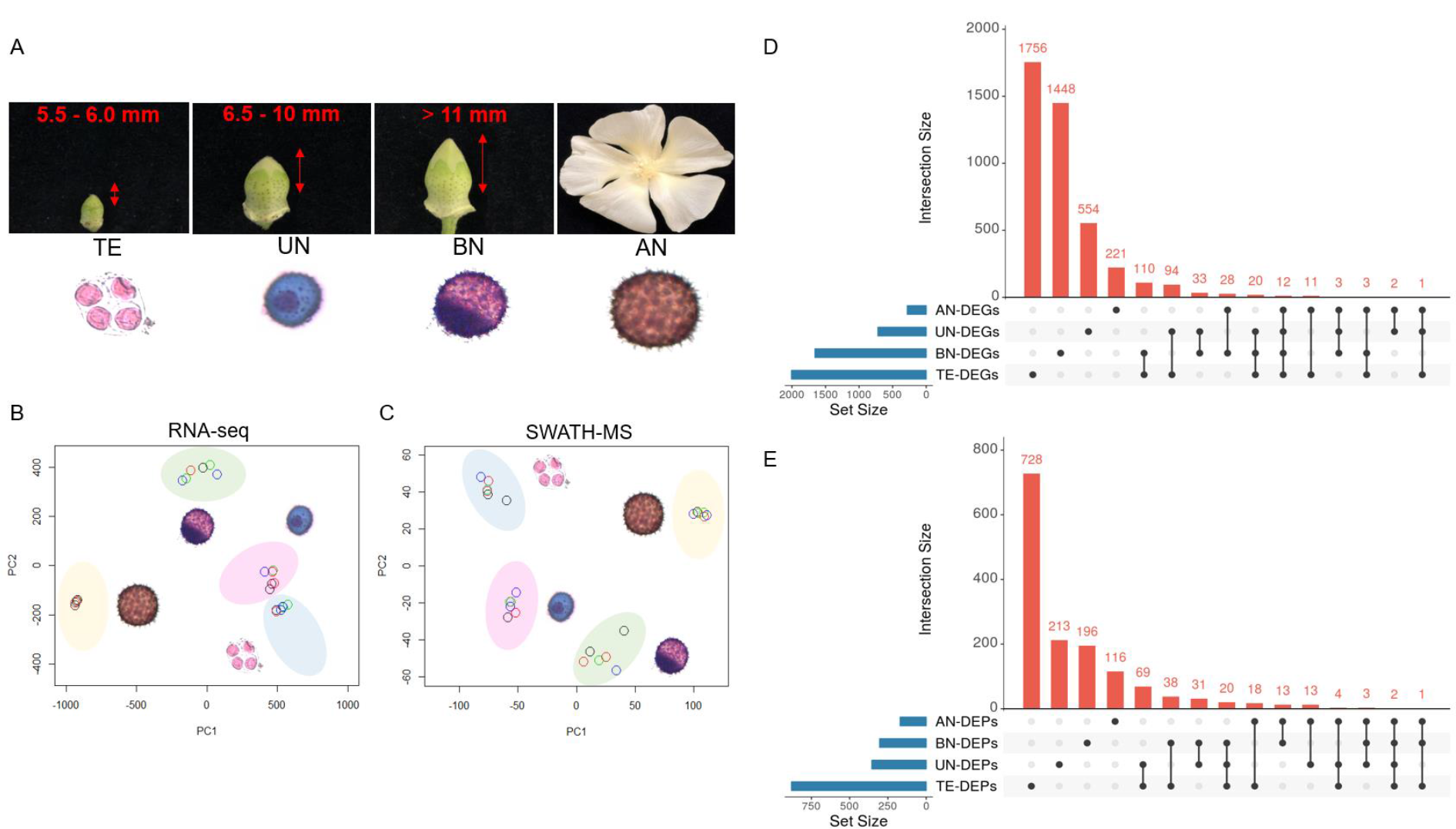
Stage-specific responses of cotton pollen developmental stages to heat stress using RNA-seq and SWATH-MS. (A) the four pollen reproductive stages including tetrads (TE), uninucleate (UN), binucleate (BN) and dehisced pollen (AN) collected for RNA-seq and SWATH-MS analysis after exposure to 38/28°C for 5 d; (B and C) principal component analysis (PCA) of total genes and proteins, respectively; (D and E) UpSet plot of DEGs and DEPs, respectively.

### RNA-seq data analysis

High-throughput sequencing was performed on 24 samples consisting of TE, UN, BN and AN under control and heat conditions with three biological replicates. The complete set of paired-end FASTQ mRNA-seq reads have been deposited at the NCBI Sequence Read Archive (SRA) with project number SUB870307. The Illumina sequencing on a NovaSeq 6000 system yielded a total of over 2,039 million paired-end 100 base reads for the 24 libraries after the sequences were trimmed. Approximately 90% of the total reads were mapped against the cotton reference genome (*Gossypium hirsutum* L. acc. TM-1), of which about 75% were mapped to genes. Gene expression level was evaluated by counting the number of mapped reads per gene, followed by TMM normalisation. The result was then reported as fold-change to indicate the changes in transcript levels at the four pollen developmental stages under the elevated temperature. Principal component analysis (PCA) on the pooled samples (total counts as entry), including four pollen developmental stages, showed that each developmental stage was distinct, demonstrating high reproducibility within groups (Fig. 2B).

### Differentially expressed genes (DEGs) in pollen developmental stages

The transcript per kilobase million (TPM) was considered to select the final set of DEGs, with the condition that at least 2 of 3 replicates must score above zero values. We identified a total of 2007, 719, 1657 and 281 genes differentially expressed in TE, UN, BN and AN, respectively, in response to heat stress (Supplementary Data Table S1). An UpSet plot was used to visualize the intersecting sets of genes, demonstrating that a high number of DEGs are stage-specific in response to heat (Fig. 2D). A total of 1,756 genes were *uniquely* differentially affected by heat in the early stage of development (TE), whereas only 221 genes were unique to the late stage (AN) being affected by exposure to 38°C. Only 12 genes were commonly changed at all stages in response to heat. The common DEGs in all stages included HSP21 (Gh_D04G1804), HSP15.7 (Gh_D11G1397), HSP18.2 (Gh_D08G1737), HSP18.5-C (Gh_D03G1268), HSP17.9-D (Gh_A05G0305), DnaJB13 (HSP40; Gh_A05G1343), HSP83A (Gh_D03G1421), FAM32A (Gh_A06G0967), HSP70 (Gh_A13G2046 and Gh_D13G2447) and HSC70 (Gh_A05G0823 and Gh_D05G0943). Transcriptome analysis showed that the highest number of up-regulated genes belonged to TE (1243), followed by UN (631), BN (584) and AN (142), whereas, in the same tissues, 764, 88, 1073 and 139 genes were down-regulated, respectively, after exposure to heat (Fig. 3A-B).

**Fig. 3.**
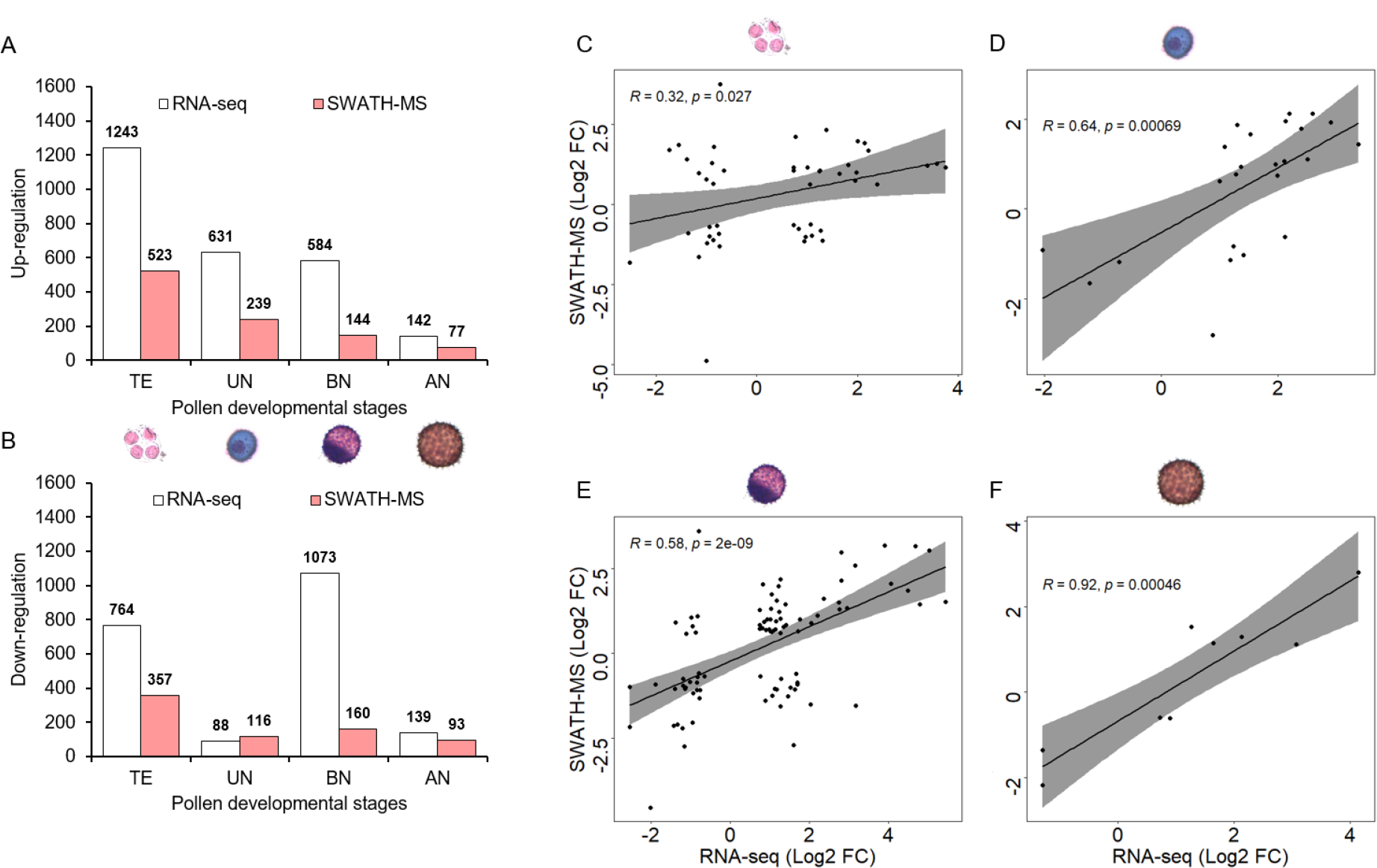
The number of DEGs and DEPs identified in pollen developmental stages using RNA-seq and SWATH-MS datasets and their correlation, after exposure of the plants to 38/28°C for 5 d. (A and B) number of up- and down-regulated genes and proteins in the four distinct reproductive stages, respectively. The relationship between correlated differentially expressed genes (DEGs) and proteins (DEPs) in TE (C), UN (D), BN (E) and AN (F). *X*-axis represents the gene expression level, and y-axis represents the protein abundance level (all data were log2-transformed).

### Generation of a spectral library for SWATH-MS

To acquire quantitative protein information of pollen development under heat stress, we generated an extensive spectral library using anther, leaf, root and pollen from cotton. From this library, we obtained reference spectra corresponding to 5,257 proteins. The spectral library was used to quantify proteins involved in TE, UN, BN and AN in response to 38°C using SWATH-MS. Proteomes of anthers (ANT) and leaves (LE) were also analysed to reveal the response of vegetative tissues to high temperature compared with the reproductive stages. This SWATH assay library is available through the ProteomeXchange Consortium via the PRIDE partner repository (Perez-Riverol et al., 2019) with the dataset identifier PXD019873, as a public resource to support further research in plant science.

### SWATH-MS data analysis

Proteomes were analysed by SWATH-MS in four pollen developmental stages (TE, UN, BN and AN), alongside ANT and LE. Using the SWATH spectral library described above, we were able to identify 10,506, 6,056 and 5,574 peptides, representing 4,501, 3,056 and 2,958 proteins in pollen, anther and leaf samples, respectively (Supplementary Data Table S2). Similar to the PCA result at transcriptome level, the total proteins (peak areas as entry) identified in the pollen developmental stages grouped separately, with all the replicates grouped together (Fig. 2C).

### Differentially expressed proteins in pollen developmental stages

Using the data obtained from SWATH-MS, we found that up to 880 proteins were differentially changed in TE after exposure to heat, with 355, 304 and 170 proteins in UN, BN and AN, respectively (Fig. 3A-B, Supplementary Data Table S3), revealing a dramatic decrease in the number of heat-responsive DEPs as pollen developed. Using an UpSet plot, 728 proteins were *uniquely* differentially regulated in TE, representing 83% of DEPs (880) identified at this stage, again illustrating the distinctive response to heat that characterised tetrads. Moreover, about two-thirds of DEPs were unique to each of the other three stages of pollen development: 213 out of 355 (60% of DEPs), 196 out of 304 (65% of DEPs) and 116 out of 170 proteins (68% of DEPs) were quantitatively changed in abundance in UN, BN and AN at 38°C, respectively. Surprisingly, only two DEPs were significantly responsive to heat at all stages analysed (Fig. 2E). The common DEPs in all stages included 15.7 kDa heat shock protein, peroxisomal-like (HSP15.7; A0A1U8L3S8) and peptidylprolyl isomerase (ROC4; A0A1U8I2M5), both of which contribute to protein refolding. Similar to RNA-seq analysis, SWATH-MS demonstrated that the four stages responded distinctively to heat, with a large number of unique genes and proteins significantly changed in the early stage of pollen development. Interestingly, we found a large number of down-regulated proteins at the late developmental stages (BN and AN) in response to heat, while the early stages (TE and UN) mostly up-regulated the proteins (Fig. 3A-B). This was also observed in the vegetative tissues where anthers and leaves down-regulated synthesis of most proteins in response to heat stress (data not shown).

### Correlation between transcriptomics and proteomics datasets

DEPs with Uniprot identifiers were blasted against the cotton reference proteome (*Gossypium hirsutum* L. acc. TM-1) to search for the corresponding cotton identifiers using a local blastp platform (OmicsBox ver 1.4.11). Although the Pearson correlation (r) was between -0.0014 and 0.16 for the total pool of genes and proteins identified (data not shown), the correlation between differentially expressed genes and proteins was high in all developmental stages, with TE having the lowest correlation (32%), followed by BN (58%), UN (64%) and AN (92%, Fig. 3C-F). The number of DEGs and DEPs varied notably in the developmental stages, and only a total of 148 correlated IDs were identified of which 49, 24, 90 and 9 IDs were associated with the early stage (TE), UN, BN and AN, respectively. It is commonly observed across the plant literature that transcript levels often correlate poorly with protein abundance, mainly because of post-transcriptional regulation. These data suggest that a high degree of regulation also occurs in gene expression after heating, with most transcripts and proteins responding independently. The minority of genes that are transcribed and translated in response to heat in a coordinated manner could be useful as markers for selective breeding programs.

### Functional analysis of pollen, anther and leaf tissues at high temperature

The most enriched gene ontology (GO) categories in DEGs are shown in Figure 4, and the most enriched GO categories for DEPs are shown in Figure 5 (Fisher’s exact test, *p*-value <0.05). A detailed list of GO enrichment analysis can be found in Supplementary Data Tables S4 and S5. At the transcript level, functional enrichment analysis demonstrated that in the early pollen stage (TE), heat-activated genes contributed to the regulation of transcription and cell wall biogenesis, whereas in the later developmental stages (UN, BN and AN), the up-regulated pathways were mainly associated with protein homeostasis, such as regulation of protein complex stability. Importantly, a substantial number of the highly enriched GOs in mature pollen (AN) were annotated as molecular chaperones, indicating the importance of protein re-folding during heat. In tetrads, important functions including pollen wall assembly, intercellular transport and response to ABA, all of which play pivotal roles in plant thermotolerance, were inhibited at high temperature. The later stages of pollen development, however, responded to heat differently in that they slowed down metabolic pathways such as xyloglucan metabolic process.

**Fig. 4.**
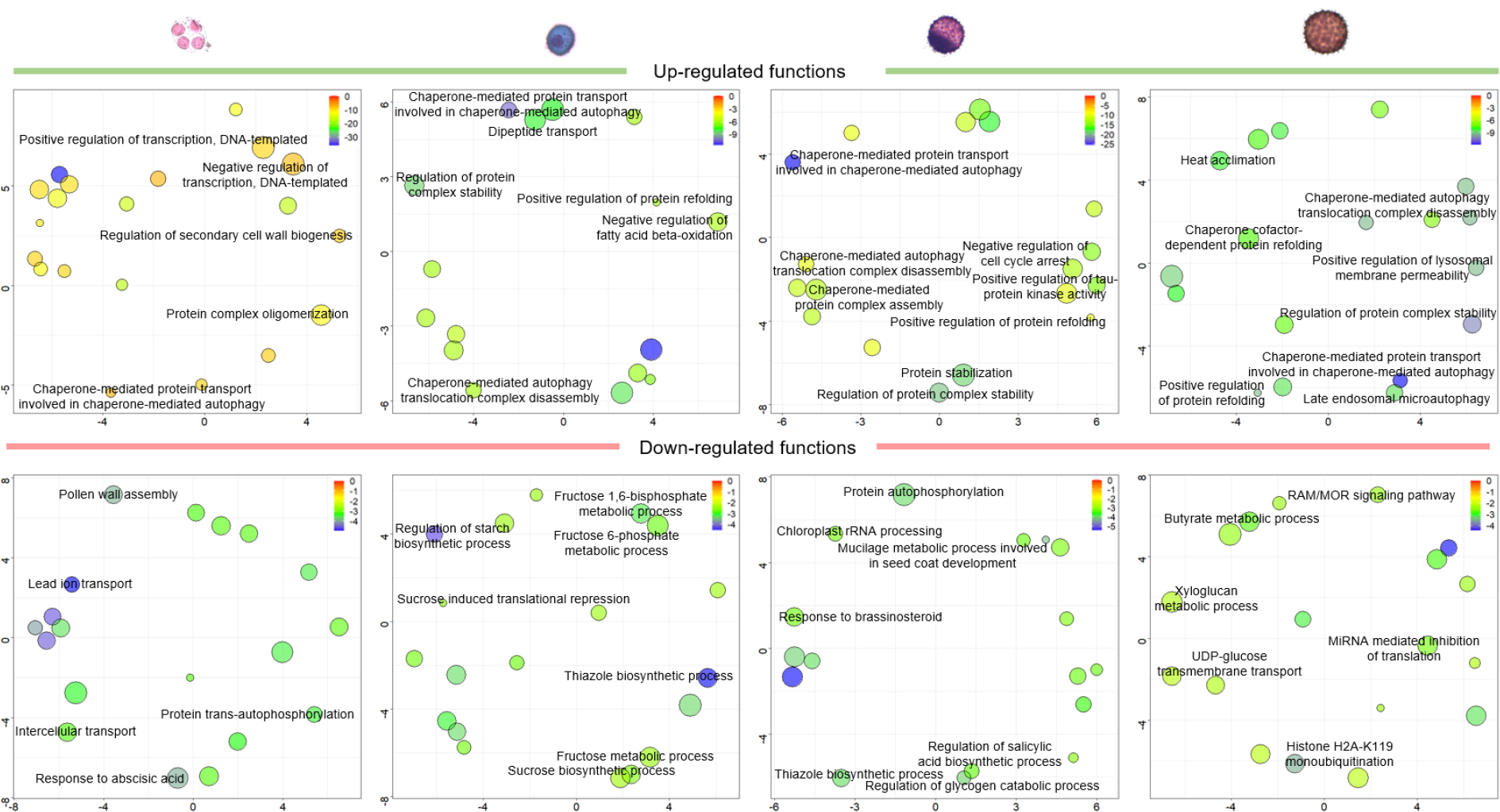
Biological processes enriched (*p*-value <0.05) in up- and down-regulated genes (DEGs), using REVIGO, in four pollen developmental stages after exposure to 5 d extreme heat. Each bubble indicates a significantly enriched term in a two-dimensional space derived by applying multidimensional scaling to a matrix of the GO terms’ semantic similarities (Supek et al., 2015). Bubble size is proportional to the frequency of the GO term in the whole Uniprot database (bubbles of more general GO terms are larger), whereas color indicates the log10 *p*-value, with red and blue representing higher and lower *p*-value, respectively. The top 20 statistically most significant GO terms are plotted, of which only the key functions were labelled.

**Fig. 5.**
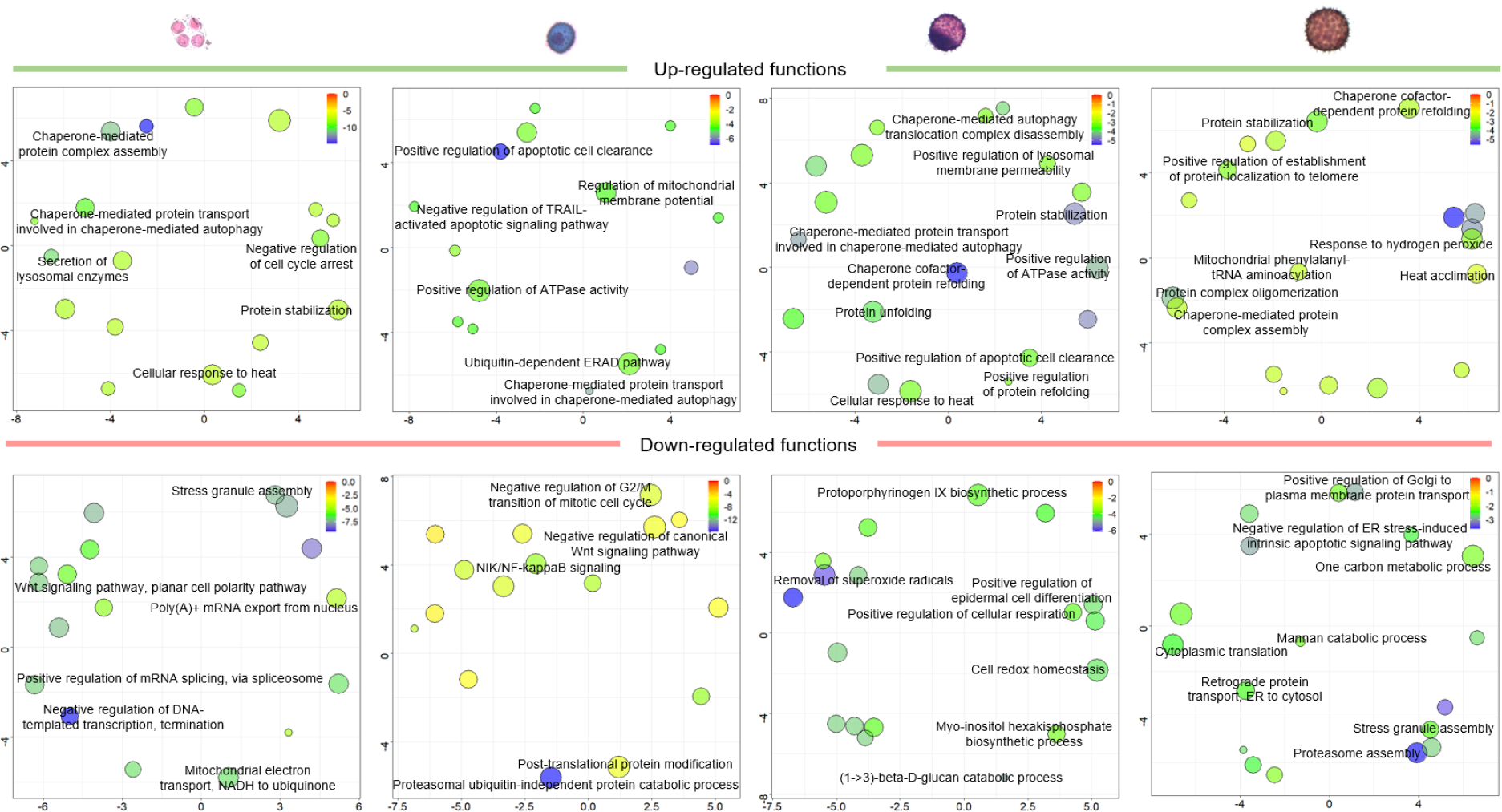
Biological processes enriched (*p*-value <0.05) in up- and down-regulated proteins (DEPs), using REVIGO, in four pollen developmental stages after exposure to 5 d extreme heat. Each bubble indicates a significantly enriched term in a two-dimensional space derived by applying multidimensional scaling to a matrix of the GO terms’ semantic similarities (Supek et al., 2015). Bubble size is proportional to the frequency of the GO term in the whole Uniprot database (bubbles of more general GO terms are larger), whereas color indicates the log10 *p*-value, with red and blue representing higher and lower *p*-value, respectively. The top 20 statistically most significant GO terms are plotted, of which only the key functions were labelled.

RNA-seq analysis demonstrated that splicing was compromised in tetrads after exposure to high temperature, as was evident from down-regulation of PRP19A, pre-mRNA-processing factor 19 homolog 1 (Gh_A01G0193), whereas two isoforms of serine/arginine-rich splicing factor SR30 (Gh_D13G1145 and Gh_A13G1537) were highly expressed in mature pollen (AN). Moreover, DEGs from mature pollen indicated that high levels of transcription could be maintained at 38°C by down-regulation of miRNA-mediated inhibition of translation and histone H2AK119 mono-ubiquitination (Figure 4). We also found that key genes contributing to pollen wall assembly were down-regulated in tetrads in response to heat, including AT-hook motif nuclear-localized protein 16 (AHL16; Gh_A09G0066, Gh_D09G0063, Gh_D11G0652 and Gh_Sca009301G01), AT-hook motif nuclear-localized protein 17 (AHL17; Gh_A02G0807 and Gh_A13G1898) and PHD finger protein MALE STERILITY 1 (MS1; Gh_D12G2265 and Gh_A12G2089). Functional enrichment analysis indicated that during the post-tetrad stages of pollen development, some metabolic processes such as starch biosynthetic process (UN), glycogen biosynthetic process (BN) and xyloglucan metabolic process (AN) were repressed in response to heat. For example, different isoforms of Cyclin-U4-1 (CYCU4-1; Gh_A11G0680, Gh_D11G0795 and Gh_D12G0972) involved in cell division were down-regulated in BN when plants were exposed to high temperature. Overall, transcriptomics indicated that the most tolerant stage of pollen development (AN) to high temperature was associated with suppression of expression of metabolic processes but maintenance of high levels of transcription.

At the protein level, GO enrichment analysis showed that all developmental stages up-regulated molecular chaperones somewhat, enabling proteins to remain functional under heat. In addition, the apoptosis pathway was activated by an increase in the level of lysosome secretion in tetrad cells, which continued up to the BN stage in response to heat. Functional analysis indicated that the cell cycle was adversely affected by exposing tetrads to extreme heat. Notably, in the tetrad stage, the signaling pathway of Wnt involved in cell division and mitochondrial electron transport were inhibited under heat stress. Importantly, transcription and mRNA splicing were inhibited in the tetrad stage after heat.

The pathway associated with response to hydrogen peroxide was activated in mature pollen, potentially reducing oxidative damage accumulated by heat (Figure 5). As observed in the transcriptomics data, during the late developmental stages, metabolic processes such as (1->3)-ß-D-glucan catabolic process, myo-inositol hexakisphosphate biosynthetic process, mannan catabolic process and the one-carbon metabolism process were down-regulated. Cytoplasmic translation was also down-regulated in mature pollen (Figure 5). Similar to AN, GO enrichment analysis of the vegetative tissues indicated that translation was down-regulated in leaves when exposed to high temperature (Figure 7A). Our finding indicated that tricarboxylic acid (TCA) cycle and gluconeogenesis were highly enriched in anthers and leaves respectively, in response to heat.

Proteomics data showed that different isoforms of glycine-rich RNA-binding protein (A0A1U8KS79, A0A1U8LME9, A0A1U8NV97, A0A1U8N4Q3, D2D310, A0A1U8K9R7, A0A1U8NZI4 and A0A1U8M9R9) and heterogeneous nuclear ribonucleoprotein 1 (RNP1; A0A1U8IAH1) involved in pre-mRNA splicing became less abundant in tetrad cells under heat stress. In contrast, only A0A1U8NV97 was down-regulated in AN after exposure to high temperature, and none of the proteins was differentially regulated in anthers and leaves. Moreover, heterogeneous nuclear ribonucleoprotein 1-like isoform X1 (A0A1U8IAH1) associated with pre-mRNA modification was down-regulated in tetrads, indicating potential failure in activity of spliceosome. DEPs demonstrated that proteins were mostly contributed to the activation of ribosomal processes in tetrads after exposure to heat. On contrary, ribosomal proteins were mostly down-regulated in mature pollen and leaves in response to high temperature, demonstrating the suppression of translational processes in the tolerant tissues. Overall, proteomics data indicated that mature pollen (AN) and leaves, as relatively tolerant tissues, translation was suppressed in response to heat stress.

### Shared GOs in pollen developmental stages differentially respond to heat

The developmentally common GO terms were plotted in polar graphs to compare the impact of extreme heat, in terms of up- or down-regulation, across the pollen developmental stages (Figure 6A-B). The analysis showed that all common terms were up-regulated in the early stage of development (TE) in response to heat, whereas down-regulation increased over the developmental stages, with the mature pollen (AN) having the largest number of GO terms down-regulated in response to heat.

**Fig. 6.**
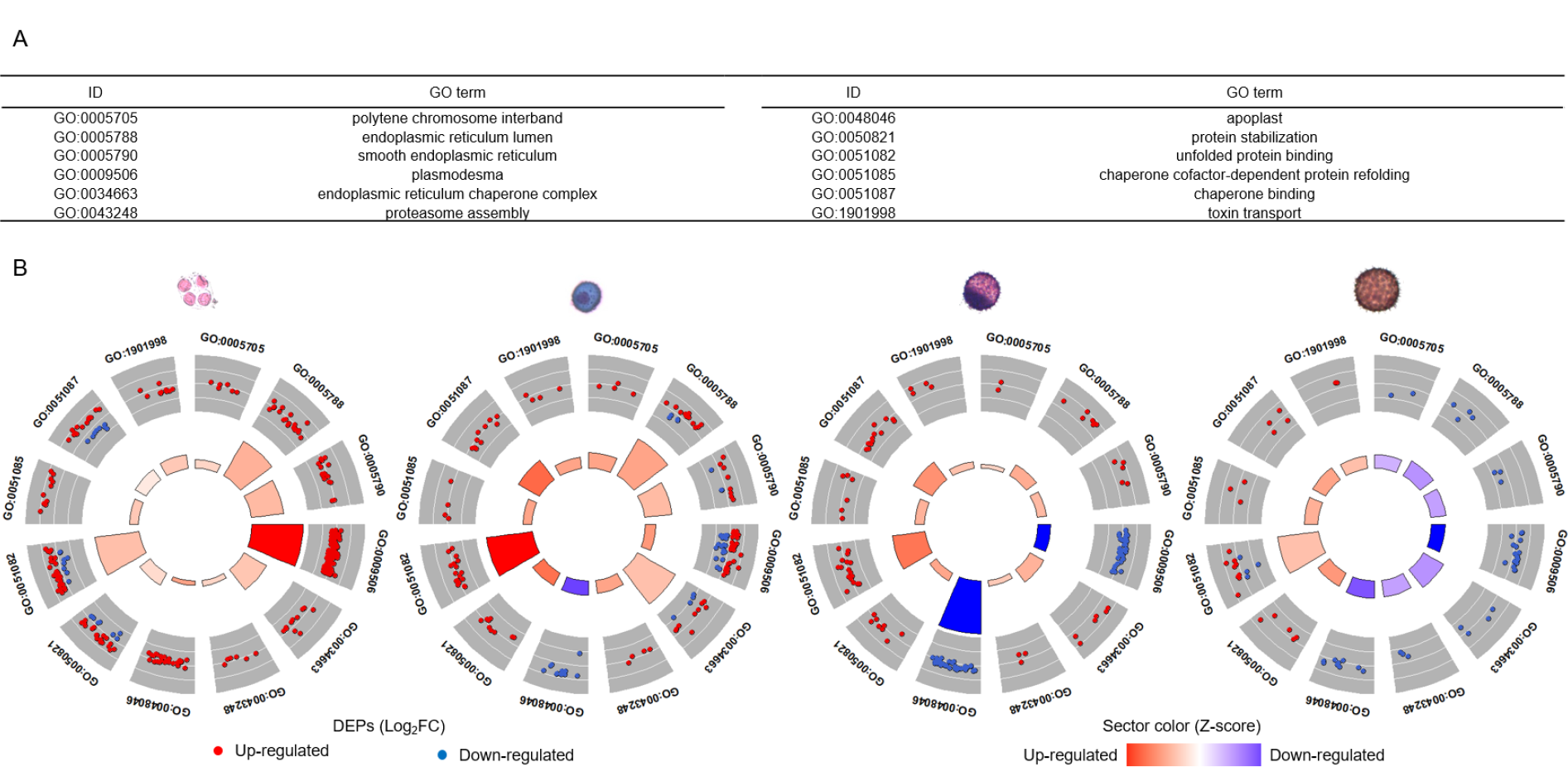
Different response of functions identified by GO enrichment analysis during pollen developmental stages at 38/28°C for 5 d. (A) the 12 identified common enriched GOs in DEPs among four pollen developmental stages including TE, UN, BN and AN; (B) polar graphs representing the direction of response of each stage to extreme heat. The outer ring shows the log 10 FC of each DEP (red and blue colours indicate up- and down-regulated proteins, respectively) annotated with a specific common GO category. The size of the sectors in the inner ring is proportional to the statistical significance (adjusted *P* value) of GOs enriched in OmicsBox. The colour shows the tendency of the GO to up- or down-regulate in response to heat according to it z score, calculated by the following formulae: 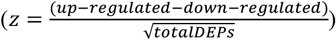.

### The vegetative tissues down-regulate the shared GO with the early reproductive stage

To further investigate the effect of down-regulation on plant thermotolerance, we analysed the vegetative and reproductive tissues because of their respective differences in thermotolerance. The shared GO terms were used to compare the response of anthers, leaves and the tetrad stage to heat. We found that anthers had up-regulated proteins in relation to the TCA and glyoxylate cycles, possibly to increase the level of carbohydrates required for pollen. Moreover, anthers up-regulated chaperones and superoxide dismutase (SOD) after exposure to heat. Proteomics data indicated that unlike pollen developmental stages and anthers, molecular chaperones were not among the most enriched functions in leaves after exposure to heat. Instead, the leaf cells activated proteins associated with gluconeogenesis and calcium ion transport from cytosol to ER to mitigate the stress. On the other hand, the apoptosis pathway was inhibited in anthers, whereas it was activated in the early pollen developmental stages in response to heat. Also, in contrast to leaves, anthers down-regulated proteins related to transport of calcium ions to the ER. Similar to pollen developmental stages, metabolic processes such as ITP catabolic process, glucose-1-phosphate metabolic process and glyceraldehyde-3-phosphate biosynthesis process, were inhibited in the vegetative tissues under heat stress. In leaves, proteins contributing to the translation processes were also down-regulated in response to heat. As a relatively heat-tolerant tissue, leaves maintained cell homeostasis by activating key pathways, and translation declined notably after exposure to heat (Figure 7A). In agreement with the results shown in Figure 6, shared GO terms among the early pollen stage (TE), anther (ANT) and leaf (LE) demonstrated that the majority of functional categories in tetrads were activated when exposed to extreme heat, while the overall tendency towards down-regulation of the same categories was greater in leaves.

The results from functional enrichment analysis indicated that, at the RNA level, transcription remained active when the later stages of pollen were exposed to heat stress. Inhibition of transcription in the tetrad stage, however, might imply the vulnerability of the tetrad stage to high temperature. Moreover, at the protein level, translation was down-regulated as seen in mature pollen and leaf when exposed to high temperature, (Figure 4-5-7A).

**Fig. 7.**
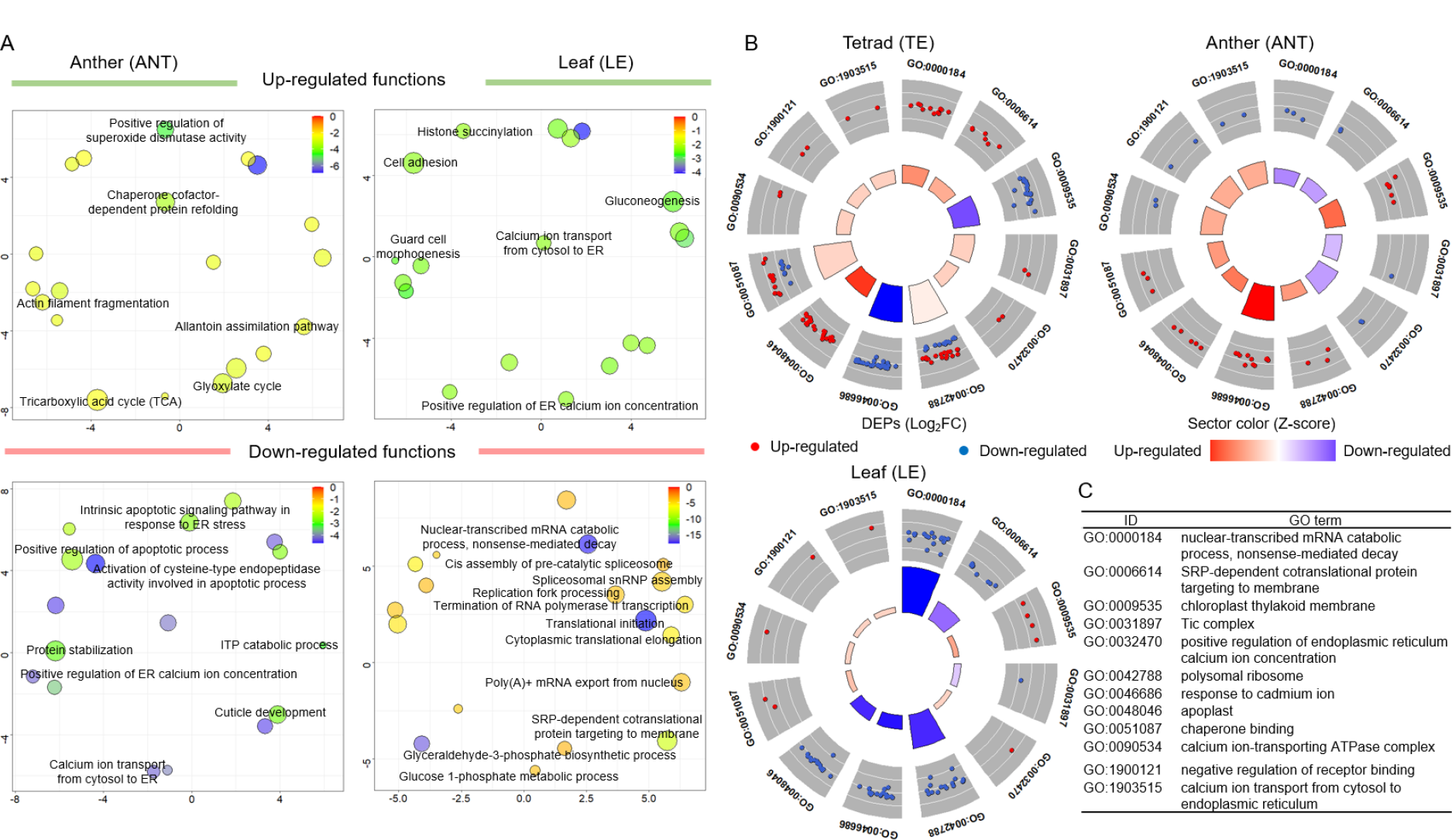
GO enrichment analysis of the vegetative tissues including anther and leaf at 38/28°C for 5 d. (A) biological processes enriched (*p*-value <0.05) in up- and down-regulated proteins (DEPs), in anther (ANT) and leaf (LE) after exposure to 5 d extreme heat. Each bubble indicates a significantly enriched term in a two dimensional space derived by applying multidimensional scaling to a matrix of the GO terms’ semantic similarities (Supek et al., 2015). Bubble size is proportional to the frequency of the GO term in the whole Uniprot database (bubbles of more general GO terms are larger), whereas color indicates the log10 *p*-value, with red and blue representing higher and lower *p*-value, respectively. The top 20 statistically most significant GO terms are plotted, of which only the key functions were labelled; (B) polar graphs representing the direction of response of each stage to extreme heat. The outer ring shows the log 10 FC of each DEP (red and blue colours indicate up- and down-regulated proteins, respectively) annotated with a specific common GO category. The size of the sectors in the inner ring is proportional to the statistical significance (adjusted *P* value) of GOs enriched in OmicsBox. The colour shows the tendency of the GO to up- or down-regulate in response to heat according to it z score, calculated by the following formulae: 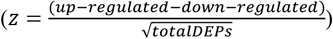; (C) 12 common enriched GOs identified in DEPs among TE, ANT and leaf.

### Protein folding was substantially activated in susceptible tissues

Clustering of the proteins differentially regulated by heat during pollen development indicated that the relative fold-change of differentially changed proteins is higher in tetrads after exposure to 38°C, in comparison with the late developmental stages (UN, BN and AN; Figure 8A). SWATH-MS data indicated that a total of 116 DEPs were associated with the pathways closely related to the folding of proteins, according to their GOs. The Heatmap (Figure 8B) demonstrates that the majority of the proteins associated with (re)folding were differentially regulated in TE, indicating a high sensitivity of tetrad cells to heat. Nevertheless, the abundance of these proteins mostly remained unchanged in the late stages of pollen development, especially mature pollen. This was more obvious in the vegetative tissues where only a few proteins related to folding were significantly changed. We found that two proteins became more abundant in all four pollen developmental stages in response to heat: 15.7 kDa heat shock protein, peroxisomal-like (A0A1U8L3S8) and peptidylprolyl isomerase (A0A1U8I2M5). DnaJ homolog subfamily B member 13-like (A0A1U8KL17) and 17.5 kDa class I heat shock protein-like (A8WCV1) were not significantly up-regulated in TE; however, their abundance increased in UN, BN and AN, after exposure to heat. Importantly, 17.3 kDa class II heat shock protein-like (A0A1U8PG98), 17.1 kDa class II heat shock protein-like (A0A1U8LTX0), T-complex protein 1 subunit zeta 1-like (A0A1U8N4I9) and T-complex protein 1 subunit eta (A0A1U8LYZ3) were only up-regulated in AN (mature pollen), which suggests they might play key roles in its tolerance to heat when compared with the immature stages of pollen. Importantly, a large number of differentially changed proteins were down-regulated in AN, ANT and LE after plants were exposed to extreme heat.

**Fig. 8.**
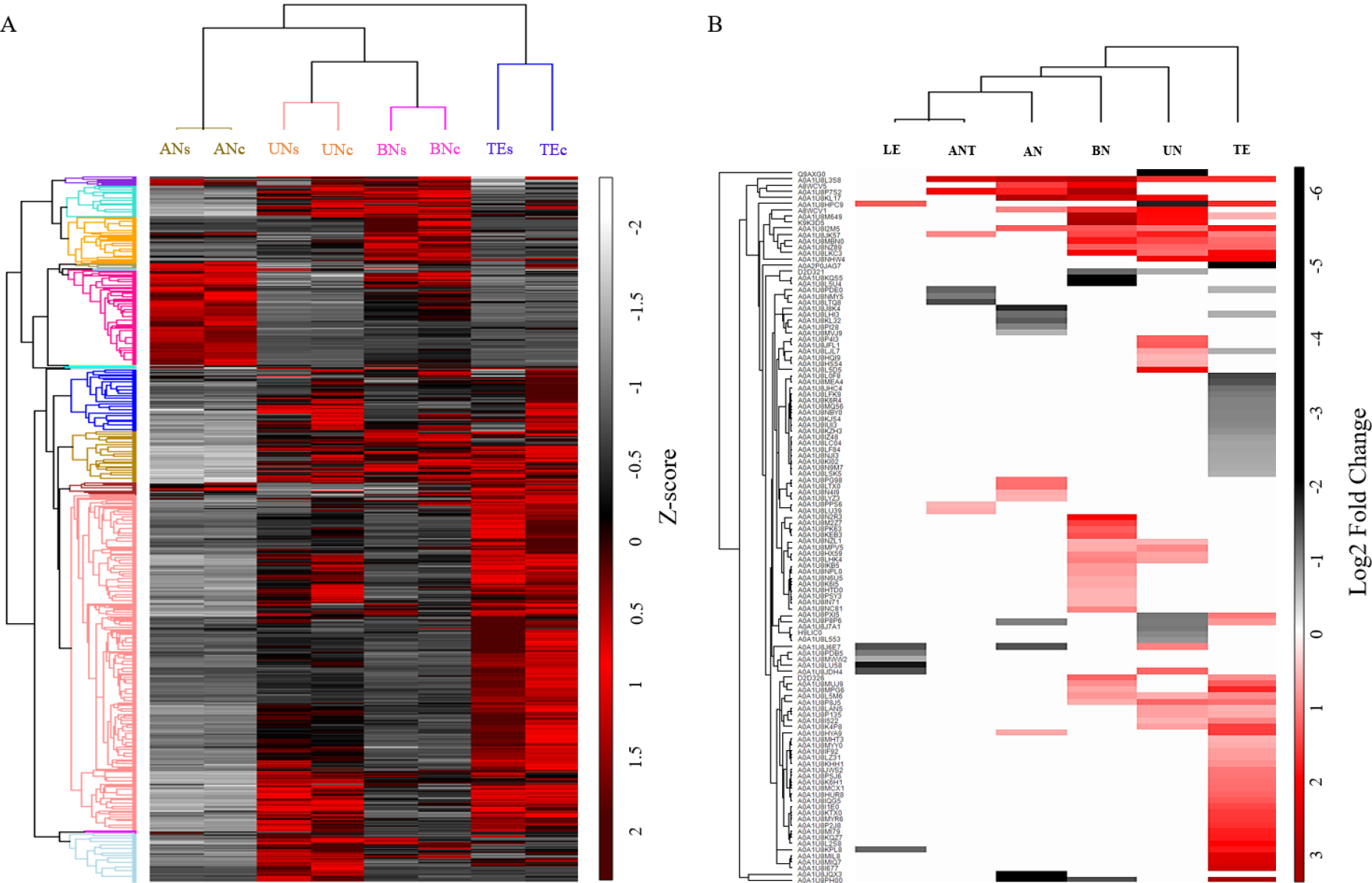
(A) hierarchical clustering of the Z-scored values of DEPs (analysis of variance (ANOVA), with *p* < 0.05 and fold-change greater than ± 1.5) created in Perseus (Version 1.6.5.0) using Euclidean as the distance metric and average as the linkage criterion (Tyanova et al., 2016); (B) expression patterns of all DEPs related to protein folding (based on their GOs) in TE, UN, BN, AN, ANT and LE involved in heat stress (38/28°C for 5 d). The fold changes of 116 significantly changed proteins were log 2-transformed and a hierarchical clustering was created in Perseus (Version 1.6.5.0) using Euclidean as the distance metric and average as the linkage criterion (Tyanova et al., 2016).

### PRM validation of heat-responsive proteins

Parallel reaction monitoring (PRM) analysis was performed on ten heat-responsive proteins to validate differentially regulated proteins obtained from SWATH-MS (Table 1). Unique peptides were selected for the proteins using Skyline and included in the PRM analysis. The results indicated that the differential changes in abundance of *protein disulfide-isomerase* (A0A1U8PXI5), *chaperonin CPN60-2, mitochondrial* (A0A1U8HX59) and *prefoldin subunit 1* (A0A1U8J7A1) agree with the SWATH-MS data in TE, UN, BN and AN. SWATH-MS was unable to identify statistically significant changes in some proteins and tissues, even though it showed the same trend (up or down), when compared with PRM. There were also a few proteins which were found to be not significant in PRM, but were significantly changed in SWATH-MS. Overall, the PRM data supported the data obtained using SWATH-MS, confirming the reliability of our proteomics datasets.

**Table 1.** Comparing PRM, SWATH-MS and RNA-seq datasets with respect to changes in fold change of several key proteins. A total of nine heat-responsive proteins including peptidylprolyl isomerase (A0A1U8I2M5), heat shock 70 kDa protein (A0A1U8MBN0), protein disulfide-isomerase (A0A1U8PXI5), chaperonin CPN60-2, mitochondrial (A0A1U8HX59), 10 kDa chaperonin-like (A0A1U8NJI3), luminal-binding protein 5-like (A0A1U8JWS2), prefoldin subunit 1 (A0A1U8J7A1), 20 kDa chaperonin, chloroplastic-like (A0A1U8LU39) and annexin (A0A1U8J8K4) were selected.

| Uniprot ID | Gh ID | TEc vs TEs |  |  | UNc vs UNs |  |  | BNc vs BNs |  |  | ANc vs ANs |  |  |
| --- | --- | --- | --- | --- | --- | --- | --- | --- | --- | --- | --- | --- | --- |
|  |  | PRM | SWATH-MS | RNA-seq | PRM | SWATH-MS | RNA-seq | PRM | SWATH-MS | RNA-seq | PRM | SWATH-MS | RNA-seq |
| A0A1U8I2M5 | Gh_D09G1979 | Up<sup>*</sup> | Up<sup>*</sup> | Up<sup>*</sup> | Up<sup>*</sup> | Up<sup>*</sup> | Up<sup>*</sup> | Up<sup>*</sup> | Up<sup>*</sup> | Up<sup>*</sup> | Up<sup>ns</sup> | Up<sup>*</sup> | Up<sup>*</sup> |
| A0A1U8MBN0 | Gh_A13G2046 | Up<sup>*</sup> | Up<sup>*</sup> | Up<sup>*</sup> | Up<sup>*</sup> | Up<sup>*</sup> | Up<sup>*</sup> | Up<sup>*</sup> | Up<sup>*</sup> | Up<sup>*</sup> | Up<sup>ns</sup> | Up<sup>ns</sup> | Up<sup>*</sup> |
| A0A1U8PXI5 | Gh_A03G0285 | Up<sup>*</sup> | Up<sup>*</sup> | Up<sup>ns</sup> | Down<sup>*</sup> | Down<sup>*</sup> | Up<sup>ns</sup> | Up<sup>ns</sup> | Up<sup>ns</sup> | Down<sup>*</sup> | Down<sup>ns</sup> | Down<sup>ns</sup> | Up<sup>ns</sup> |
| A0A1U8HX59 | Gh_D09G1918 | Up<sup>ns</sup> | Up<sup>ns</sup> | Up<sup>ns</sup> | Up<sup>*</sup> | Up<sup>*</sup> | Up<sup>*</sup> | Up<sup>*</sup> | Up<sup>*</sup> | Up<sup>*</sup> | Up<sup>ns</sup> | Up<sup>*</sup> | Down<sup>ns</sup> |
| A0A1U8NJI3 | Gh_A08G0086 | Down<sup>*</sup> | Down<sup>*</sup> | Up<sup>*</sup> | Down<sup>ns</sup> | Down<sup>ns</sup> | Up<sup>ns</sup> | Up<sup>ns</sup> | Up<sup>ns</sup> | Up<sup>ns</sup> | Down<sup>ns</sup> | Up<sup>ns</sup> | Down<sup>*</sup> |
| A0A1U8JWS2 | Gh_D08G1192 | Up<sup>*</sup> | Up<sup>*</sup> | Down<sup>ns</sup> | Up<sup>*</sup> | Up<sup>ns</sup> | Up<sup>*</sup> | Up<sup>ns</sup> | Up<sup>ns</sup> | Up<sup>*</sup> | Down<sup>ns</sup> | Up<sup>ns</sup> | Down<sup>ns</sup> |
| A0A1U8J7A1 | Gh_A05G2453 | Down<sup>ns</sup> | Down<sup>ns</sup> | Up<sup>ns</sup> | Down<sup>*</sup> | Down<sup>*</sup> | Down<sup>ns</sup> | Up<sup>ns</sup> | Up<sup>ns</sup> | Down<sup>ns</sup> | Up<sup>ns</sup> | Up<sup>ns</sup> | Up<sup>ns</sup> |
| A0A1U8LU39 | Gh_A07G2354 | Down<sup>ns</sup> | Down<sup>ns</sup> | Up<sup>ns</sup> | Up<sup>ns</sup> | Up<sup>*</sup> | Up<sup>*</sup> | Up<sup>ns</sup> | Up<sup>ns</sup> | Up<sup>*</sup> | Down<sup>ns</sup> | Up<sup>ns</sup> | Up<sup>*</sup> |
| A0A1U8J8K4 | Gh_D05G2674 | Down<sup>ns</sup> | Up<sup>ns</sup> | Down<sup>*</sup> | Down<sup>ns</sup> | Up<sup>ns</sup> | Down<sup>ns</sup> | Down<sup>ns</sup> | Up<sup>ns</sup> | Up<sup>ns</sup> | Down<sup>*</sup> | Down<sup>*</sup> | Down<sup>ns</sup> |

\* and <sup>ns</sup> show significant and non-significant differences, respectively, between control and heat-stressed samples indicated by *t*-test analysis for *p*-value < 0.05.

## DISCUSSION

Previous studies have reported a 80 - 100% reduction in fruit set after exposure of male reproductive tissues to heat; damaging heat stress is reported in maize (40°C), sorghum (39°C), pearl millet (40°C) and tomato (32°C) (Jagadish, 2020). Cotton yield was also reduced by 65% when pollen was exposed to 40°C for 5 d (Masoomi-Aladizgeh et al., 2020). We previously confirmed tetrads as the most vulnerable stage of pollen development to heat in a commercial cotton cultivar. To reveal details of the underlying molecular events after short periods of damaging but sub-lethal heat (38°C for 5 d), we perfected the collection and analysis of samples of (at most) 20 mg at four stages of pollen development (TE, UN, BN and AN) using transcriptomics and proteomics technologies. Anthers and leaves were also analysed and compared with the proteomic response of reproductive tissues after high-temperature treatment.

### Lower transcription and higher translation in tetrads in response to high temperature

Cui and Xiong (2015) stated that abiotic stress increases the need for expression of specific stress-responsive genes; however, inadequate splicing machinery leads to inaccurately spliced mRNA. For instance, it was shown that pre-mature HSP70 was highly accumulated at 45°C in maize pollen compared with 40°C, indicating a potential splicing defect for this protein under severe heat (Hopf et al., 1992). In agreement with the above-mentioned studies, RNA-seq indicated that transcription was negatively affected in the tetrad cells in response to heat, whereas it was maintained at a high level in mature pollen. Maintenance of transcription in AN is consistent with down-regulation of miRNA-mediated inhibition of translation and histone H2AK119 mono-ubiquitination GO categories (Figure 4). Improving the efficacy of splicing machinery may increase the thermotolerance of tetrads in cotton.

Contrary to rates of transcription, functional enrichment analysis of proteomics data revealed that translation was strongly inhibited in mature pollen (AN) when exposed to high temperature. For instance, eukaryotic translation initiation factors (eIFs) were only up-regulated in tetrad cells after exposure to heat, while these proteins were down-regulated in UN, BN, AN and particularly leaves. Moreover, the shared GOs highlighted that most of the shared functional categories were down-regulated in leaves exposed to heat, compared with the tetrad cells (Figure 7B). Likewise, the polar plots shown in Figure 6B reveal that common terms were up-regulated in tetrads, whereas a trend towards down-regulation increased over the developmental stages in response to heat. We postulate that for developing pollen cells to become thermotolerant, acclimation requires suppression of translation to conserve energy for survival, analogous to the *quiescence* phenomenon in the stress-biology literature. For example, Voesenek and Bailey-Serres (2013) noted that flood-tolerant plants acclimate to inundation through quiescence, conserving energy and carbohydrates for later growth. Moreover, a study on desiccation-tolerant and -intolerant algae revealed that all taxa up-regulated critical protective genes, whereas the tolerant taxon down-regulated cellular metabolism during water stress to conserve energy (Peredo and Cardon, 2020). Those genes involved in slowing down metabolism at the AN and LE stages under heat are likely to confer thermotolerance; these candidates should be further investigated as the basis of thermotolerance traits in susceptible species and tissues.

### Molecular chaperones are highly up-regulated in tetrads

Heat shock proteins (HSPs), as molecular chaperones, are responsible for protein folding, assembly, translocation and degradation under normal conditions, as well as protein refolding under stress conditions (Wang et al., 2004). It is widely accepted that plants confer tolerance to abiotic stresses by over-expression of HSPs. Sable et al., (2018) reported that inhibition of *HSP70* and *HSP90* accumulated oxidative stress, which consequently led to autophagy of the cotton ovule. Similar to earlier studies, our transcriptomics data indicated that the up-regulated genes were significantly enriched for molecular chaperones in pollen developmental stages, after exposure to 38°C. The molecular chaperones accounted for the most enriched GOs in the later developmental stages compared with TE, where these cells were associated with secondary cell wall biogenesis and transcriptional regulation (Figure 4). Moreover, proteomics data also confirmed that the majority of the up-regulated proteins were mostly enriched for molecular chaperones (Figure 5). This suggests that for pollen to acquire thermotolerance, HSPs should be up-regulated at both transcriptome and proteome levels. Even though these chaperones are required for normal cell function, insufficient levels may lead to accumulation of misfolded proteins and cellular damage when heat stress challenges rapidly developing pollen cells. On the other hand, molecular chaperones were not among the most enriched GOs in leaves after exposure to high temperature (Figure 7). This is consistent with our observation that an ambient temperature of 38°C did not damage leaves, which were tolerant to temperatures up to 40°C. This would account for the UPR pathway not being triggered to activate synthesis of molecular chaperones.

### Transport increased in tetrads after exposure to heat

Intercellular signaling processes in plant cells occur via plasmodesmata or apoplasts, which facilitate the communication between adjacent cells by controlling the transport of molecules. Identifying proteins associated with these structures would help us understand how molecular transport is regulated by plant development and environmental stimulus (Stahl and Simon, 2013; Benitez-Alfonso, 2014; Roberts and Oparka, 2003). It is evident that cellular transport is regulated by calcium, and that accumulation of ROS due to environmental stresses leads to changes in calcium fluxes in cells (Suzuki et al., 2011; Baluška et al., 2001). For instance, Holdaway-Clarke et al., (2000) demonstrated that a slight increase in cytoplasmic calcium in maize in response to stress resulted in immediate closure of plasmodesmata. Functional analysis highlighted that all proteins associated with plasmodesma (GO:0009506) and apoplast (GO:0048046) were up-regulated in the tetrad cells, after exposure to 38°C (Figure 6B). However, the corresponding enriched proteins in these categories were mainly down-regulated in later stages, particularly in BN and AN. Moreover, GO enrichment analysis of reproductive and vegetative tissues showed that enriched apoplastic proteins were down-regulated in leaf tissues in response to heat. This implies that the most vulnerable cells (tetrads) require enhanced levels of transport and signaling to survive high temperatures, whereas later stages of pollen development and leaf tissues, which are relatively heat tolerant, suppressed communication between cells as part of some as-yet undefined stress response mechanism. Consistent with our results, it was shown that osmotic stress increased the plasmodesmal permeability in plant cells, which might be due to the high requirement for solutes in damaged cells (Roberts and Oparka, 2003), and by analogy, tetrads.

### Lack of plasticity in tetrads to respond to high temperature

The ability to overcome the damaging effects of high temperature has enabled plants to colonise extreme environments such as hot deserts. Previous studies have established that refolding aggregated proteins mediated by HSPs is one of the adaptive strategies in plants to overcome the adverse effects of heat (Wahid et al., 2007; Ul Haq et al., 2019). As well as this role of HSPs in refolding of denatured proteins, a recent study indicated that protection of cells from high temperature also requires degradation of denatured proteins (X.M. Li et al., 2015). This relates to the conservative strategy in plants that appears to be central to abiotic stress tolerance, where down-regulation of non-essential genes avails cells of the resources needed to express key genes in response to heat (X. Li et al., 2015). A large number of HSPs in the early stage of pollen development (TE) reflects the susceptibility of the cells to high temperature, and consequently more HSPs were synthesized to refold the proteins damaged by heat stress (Figure 8B). On the other hand, these proteins were less up-regulated in the relatively heat-tolerant stages that follow (UN, BN and AN), and even more particularly in anther and leaf tissues. This indicates that in the most vulnerable stages of pollen development, cells invest more energy recovering denatured proteins after stress by up-regulating pathways such as UPR. However, maturing pollen, anthers and leaves are relatively heat-tolerant tissues and therefore, did not strongly activate HSPs in response to heat. In these differentiated tissues, we speculate that misfolded proteins were preferentially degraded rather than being refolded via a UPR pathway. It seems that suppression of non-essential genes, and therefore saving resources is a key adaptive approach for plants to combat unfavourable conditions.

## CONCLUSION

It is well-established that the early stage of pollen development (meiosis or post-meiosis) in almost all flowering plants is highly vulnerable to changing environments, particularly short periods of heat. However, the molecular responses of tetrad cells exposed to heat must be understood in order to develop new heat-tolerant varieties. Using RNA-seq and SWATH-MS approaches, we found that transcription was adversely affected by heat, whereas translation generally increased in tetrads. However, the opposite pattern was observed in mature pollen and leaves, being tolerant tissues, indicating the capacity to maintain expression of stress-responsive genes and suppression of non-essential translational processes to conserve resources under heat stress. Our results demonstrated that even though HSPs are required to refold the aggregated proteins under heat, molecular chaperones are not necessarily used to acclimate to extreme heat in thermotolerant tissues. This might be an adaptive strategy for plants to suppress those cellular processes that consume most energy, and that refolding denatured proteins to restore normal activity needs more energy than their degradation. Moreover, the data indicated that transport was highly activated in tetrad cells in response to high temperature, while in the later stages of pollen development and leaf tissues, dependence on cell-to-cell communication decreased. This can be associated with the down-regulation of translational processes in tolerant tissues to respond to heat efficiently. Overall, we conclude that thermotolerance in tetrads would require improved splicing machinery through over-expression of key splicing genes. Importantly, translation of non-essential proteins should be repressed so that tetrad cells can expend the scarce resources synthesizing heat-responsive proteins.

## ACKNOWLEDGMENTS

F.M.-A. acknowledges support from Macquarie University in the form of the iMQRES, MQRES and PGRF scholarships. F.M.-A. acknowledges support from BioBam team (OmicsBox) in the form of student scholarship for functional analysis of genes and proteins. F.M.-A. acknowledges use of the Plant Growth Facility (PGF), Australian Proteome Analysis Facility (APAF) and Macquarie University Faculty of Science and Engineering Microscope Facility (MQFoSE MF) and their valuable support. F.M.-A. and B.J.A acknowledge Bioplatforms Australia for support in conducting transcriptomics analysis.

## AUTHORS’ CONTRIBUTION

F.M.-A. and B.J.A. designed and developed the research. F.M.-A. cultivated and treated the plants, collected samples, performed the proteomics and transcriptomics experiments, interpreted the results and prepared the original draft of the manuscript. M.J.M. assisted with SWATH-MS analysis and interpretation of the proteomic data. Y.A. assisted with transcriptomics analysis and R scripting for data visualization. P.A.H and B.J.A. contributed to data interpretation and preparation of the final version of manuscript. All contributed to revision of the manuscript.

## DATA AVAILABILITY STATEMENT

The proteomics data were deposited in PRIDE (https://www.ebi.ac.uk/pride) under the accession number PXD019873. RNA-seq data are available in the NCBI Sequence Read Archive (SRA) at https://www.ncbi.nlm.nih.gov/sra under the accession number SUB8703075. These data will be available publicly after the date of publication.

